# Mitochondrial genome annotation and phylogenetic placement of *Oreochromis andersonii* and *O. macrochir* among the cichlids of southern Africa

**DOI:** 10.1101/393660

**Authors:** Ian Bbole, Jin-Liang Zhao, Shou-Jie Tang, Cyprian Katongo

**Affiliations:** Department of Fisheries, Mansa, Zambia; Key Laboratory of Freshwater Aquatic Genetic Resources, Ministry of Agriculture, Shanghai Ocean University, Shanghai 201306, China; Centre for Research on Environmental Ecology and Fish Nutrition (CREEFN), Ministry of Agriculture, Shanghai Ocean University, Shanghai 201306, China; National Demonstration centre for Experimental Fisheries Science Education, Shanghai Ocean University, Shanghai 201306, China; Biological Sciences Department, University of Zambia, Lusaka, Zambia

## Abstract

Genetic characterization of southern African cichlids has not received much attention. Here, we describe the mitogenome sequences and phylogenetic positioning of *Oreochromis andersonii* and *O. macrochir* among the cichlids of southern Africa. The complete mitochondrial DNA sequences were determined for *O. andersonii* and *O. macrochir*, two important aquaculture and fisheries species endemic to southern Africa. The complete mitogenome sequence lengths were 16642 bp and 16644 bp for *O. andersonii* and *O. macrochir* respectively. The general structural organization follows that of other teleost species with 13 protein–coding genes, 2 *rRNAs*, 22 *tRNAs* and a non-coding control region. Phylogenetic placement of the two species among other African cichlids was performed using Maximum Likelihood (ML) and Bayesian Markov-Chain-Monte-Carlo (MCMC). The consensus trees confirmed the relative positions of the two cichlid species with *O. andersonii* being very closely related to *O. mossambicus* and *O. macrochir* showing a close relation to both species. Among the 13 mitochondrial DNA protein coding genes *ND6* may have evolved more rapidly and *COIII* was the most conserved. There are signs that *ND6* may have been subjected to positive selection in order for these cichlid lineages to diversity and adapt to new environments. More work is needed to characterize the southern Africa cichlids as they are important species for capture fisheries, aquaculture development and understanding biogeographic history of African cichlids. Bioconservation of some endangered cichlids is also essential due to the threat by invasive species.

## Introduction

Africa is the origin centre for cichlid diversity with well over 2000 species having diverse morphology, behaviour and ecology [1]. In Southern Africa *Oreochromis andersonii* (Castelnau 1861) and *Oreochromis macrochir* (Boulenger 1912) are two important mouth brooding endemic cichlid species in this region [2]. The former occurs in the upper Zambezi, Middle Zambezi, Kafue, Okavango and Cunene Rivers while the latter is distributed in the Upper Zambezi, Kafue, and Congo River systems [2-4]. *Oreochromis macrochir* has further been introduced to the Hawaiian Islands, Okavango and Ngami region and Cunene River basin [4]. *Oreochromis andersonii* (Three-spot tilapia) and *O. macrochir* (Green head tilapia) are important for both capture fisheries and aquaculture in Southern Africa [5]. However, due to the increase in fishing pressure as a result of an ever growing human population in this region as well as the introduction of Nile tilapia (O. *niloticus*) in almost all river systems where the native species occur, populations of these native species has greatly dwindled to vulnerable levels [6,7]. Nile tilapia hybridization with these native species and consequent decline in their population may as well make these species critically endangered in some southern African rivers such as the Kafue River [8-10]. Despite Nile tilapia dominating aquaculture production in Southern Africa, efforts are been made to domesticate native species. This is partly as a result of the growing concern of ecosystem changes in most river systems due to Nile tilapia invasion. A number of studies in Zambia have shown potential for aquaculture of some native tilapia species [11-14]. Infact, the Zambian Department of Fisheries have adopted *O. andersonii* as a candidate species for aquaculture development [15].

Molecular genetic studies on cichlids in Africa have been biased towards phylogenetics of East African Lakes [16-18]. Nile tilapia has also received global attention because of its importance as an edible fish [19]. A few peer reviewed scientific papers have reported the use of molecular genetics in the study of southern African cichlids. Phylogenetic relationships have been inferred among and between cichlid species based on selected mitochondrial DNA regions and some allozymes [20-24]. However, none of these phylogenetic analyses were based on complete mitogenome sequences.

Complete mitogenome sequences have been employed in phylogenetic analyses of different fish species [25-33]. Among the important cichlids for aquaculture in Southern Africa *(O. andersonii, O. macrochir*, *Tilapia rendalli*, *O. mossambicus* and *O. niloticus*) complete mitogenome sequence have only been determined for *O. mossambicus* and *O. niloticus*.

In this study we describe the complete mitogenome sequences of *O. andersonii* and *O. macrochir* deposited in the GenBank with accession numbers MG603674 and MG603675 respectively. Based on the mitogenome sequences of these two species and other cichlids we confirm the position of these species among the cichlid species of Africa and analyse the evolutionary rates of the protein coding genes.

## Materials and methods

### Sample collection and DNA extraction

Tissue samples (fin clips) of *O. andersonii* and *O. macrochir* were collected from Upper Zambezi River (16.10 S, 23.295 E) and Lake Bangweulu (11.35 S, 29.58 E) in Zambia respectively with approval from the Zambian Department of Fisheries. These areas had no report of *O. niloticus* presence at the time of collecting the tissues. DNA was extracted from fin clips using a TIANamp Marine Animals DNA Kit using the manufacturer’s instructions (Tiangen, China). All applicable international guidelines for the care and use of animals were followed.

### PCR amplification and sequencing

Thirty primers were designed for both *O. andersonii* and *O. macrochir* and 2 other primers for each species (S1Table). The primers were designed using aligned complete mitogenomes of *O. mossambicus* (Accession number: AY597335.1) and *O. niloticus* (Accession number: GU370126.1) [30]. PCR was performed using an Eppendorf Thermal Cycler (Eppendorf, Germany). The total reaction mixture of 25.0 μl containing 17.5 μl distilled water, 4.5 μl PCR mix (Tiangen, China), 1.0 μl forward primer, 1.0 μl reverse primer, and 1.0 μl template DNA (50 ng/ μl) was used. The reaction was denatured at 94°C for 5 minutes, followed by 35 cycles of denaturation at 94 °C for 30 seconds, annealing at 48-53°C for 30 seconds and elongation at 72°C for 45 seconds; the last extension step was carried out at 72°C for 7 minutes. Agarose (1.0%) electrophoresis was performed to visualize the PCR products. The PCR products were purified using the 3S Spin PCR product Purification Kit (Biocolor Inc., Shanghai, China). The purified DNA was sequenced on an ABI 3730 xl capillary sequencer employing the same primers as for PCR. The various regions of mitochondrial DNA were sequenced and projected as electronic outputs by the computer connected to the sequencer.

### Sequence editing, alignment and annotation

The raw sequences of the various regions of mitochondrial DNA of *O. andersonii* and *O. macrochir* were edited and assembled using BioEdit Version 7.2.6 [34]. They were further edited and aligned using complete mitogenomes of *O. niloticus* and *O. aureus* [30], *O. variabilis* [32] and *O. mossambicus* (Assession number: AY597335.1). Blast database searches on NCBI site were performed to verify the target sequences amplified. Transfer RNA *(tRNA*) genes and their secondary structures were identified using tRNAScan-SE 2.0 [35]. MEGA Version 7.0.26 [36] was used to calculate the composition of amino acids, nucleotides and codon usage in the sequences. Annotation of the sequences was performed with DOGMA [37], MITOS [38], and MitoAnnotator [39]. Further verification was done with *O. niloticus* and *O. mossambicus* genome organization. The nucleotide composition skewness was measured following the formulas: AT skew [(A – T)/(A + T)] and GC skew [(G – C)/(G + C)] [40]. MEGA Version 7.0.26 was used to test for mode of selection in protein coding genes acting at non-synonymous sites using the ration *dN/dS*.

### Phylogenetic analysis

To infer phylogeny, 29 other cichlids and another 18 non-cichlid species from 7 different families mainly found in southern Africa were obtained from the NCBI site (Table 1). The protein-coding genes were used for phylogenetic analysis except *ND6* which is encoded by the opposite strand and considered to possess a distinct heterogeneous base composition than the other 12 protein-coding genes [41]. The concatenated protein sequences were aligned using MUSCLE with default settings [42].

**Table 1.**
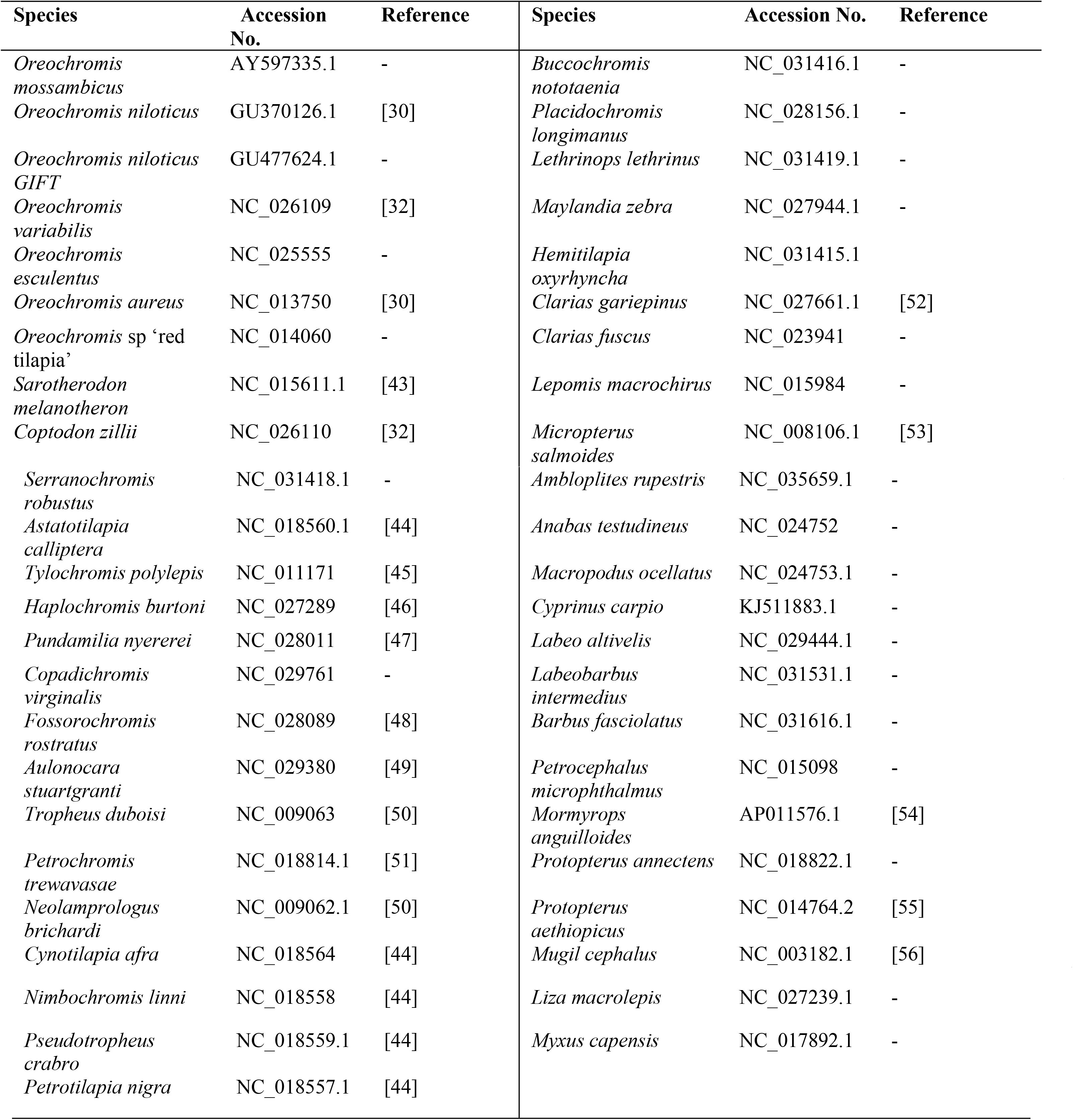
List of species with their accession numbers used in phylogenetic analysis in this study.

Phylogenetic analysis was performed using Maximum Likelihood (ML) in MEGA Version 7.0.26 [36] and Bayesian Markov-Chain-Monte-Carlo (MCMC) method in MrBayes (Version 3.2.6) [57]. Maximum Likelihood method was based on the General Time Reversible model. A discrete Gamma distribution was used to model evolutionary rate differences among sites (5 categories (+G, parameter = 0.4073)) and it allowed for some sites to be evolutionarily invariable ([+I], 15.08% sites). The bootstrap consensus tree was inferred from 1000 replicates taken to represent the evolutionary history of the taxa analyzed. Branches corresponding to partitions reproduced in less than 50% bootstrap replicates were collapsed. The Bayesian posterior probabilities were estimated using 1,000,000 generations, sampling every 100 generations at which point the standard deviation of split frequencies of the two independent runs was below 0.01. A consensus tree was constructed from the saved trees after discarding the first 25% trees as burn-in.

## Results and Discussion

### Mitogenome annotation

Annotation results of *O. andersonii* and *O. macrochir* using three methods; DOGMA, MTOS and MitoAnnotator [37-39] revealed some differences in sizes of the protein coding genes and the two *rRNAs*. However, annotation results using the three methods were similar for all *tRNAs* (S2 Table). The largest difference was observed in gene *ND5* were DOGMA and MITOS differed from MitoAnnotator in the final position with 495 bp gene size. The gene sizes for *O. andersonii* and *O. macrochir* were similar in all the three annotation methods used except *ND2* final position with a difference of 7 bp and *ND5* initial position with a difference of 3 bp (S2 Table). Comparing the results obtained, annotation from Mitoannotator was selected for submission to the Genbank (Table 2).

**Table 2.**
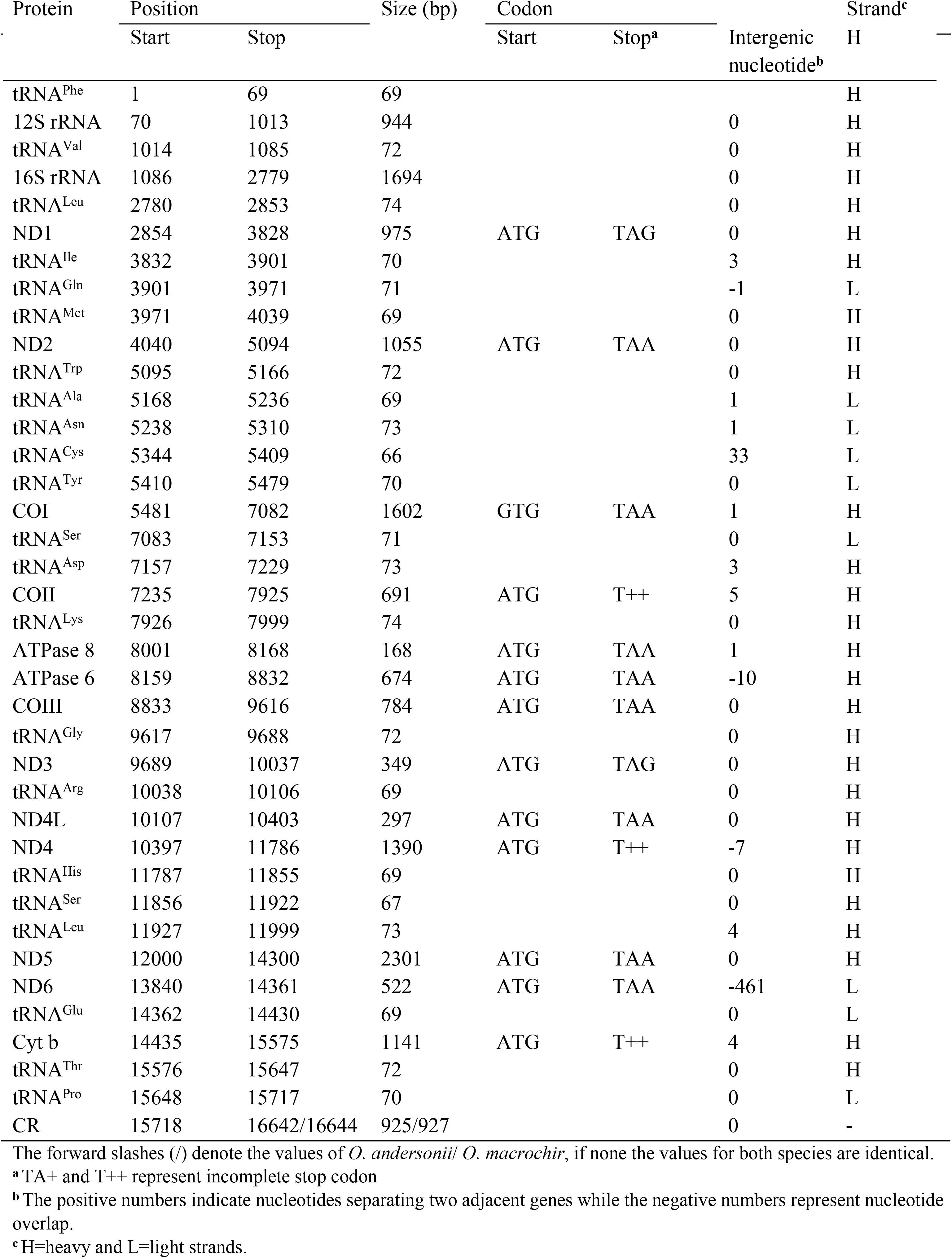
Complete automated annotated sequences of *O. andersonii* (16642 bp) and *O. macrochir* (16644 bp) generated in MitoAnnotator

### Mitogenome organization

The complete mitogenome sequences of *O. andersonii* and *O. macrochir* were 16642 bp (accession #: MG603674) and 16644 bp (accession #: MG603675) respectively. This was within the range of published mitogenomes sequences of *Oreochromis* species i.e *O. niloticus* with 16625 bp and *O. aureus* with 16628 bp [30]; *Oreochromis variabilis* with 16626 bp [32] and GenBank deposited sequence of *O. mossambicus* (AY597335.1). As anticipated, the general structural organization follows those of other teleost species with 13 protein coding genes, 2 *rRNAs*, 22 *tRNAs*, and a noncoding control region (Table 2 and figure 1). Both species were observed to exhibit the Heavy (H) and Light (L) strand coding pattern previously observed in other teleosts. The L-strand was observed in one protein coding gene *(NADH* dehydrogenase subunit 6 *(ND6)*) and eight *tRNAs (tRNA^Gln^, tRNA^Ala^, tRNA^Asn^, tRNA^Cys^, tRNA^Tyr^, tRNA^Ser^, tRNA^GIu^, tRNA^Pro^*). The start codons for protein coding genes for both *O. andersonii* and *O. macrochir* were ATG except for gene *CO1* which was GTG. The stop codons on the other hand were either TAG or TAA except for gene *CO11, ND4*, and *Cyt b* which had incomplete stop codons of T++. The incomplete codon stops observed for the two species is a common feature in most vertebrates including some fish species [27, 28, 33].

**Fig. 1.**
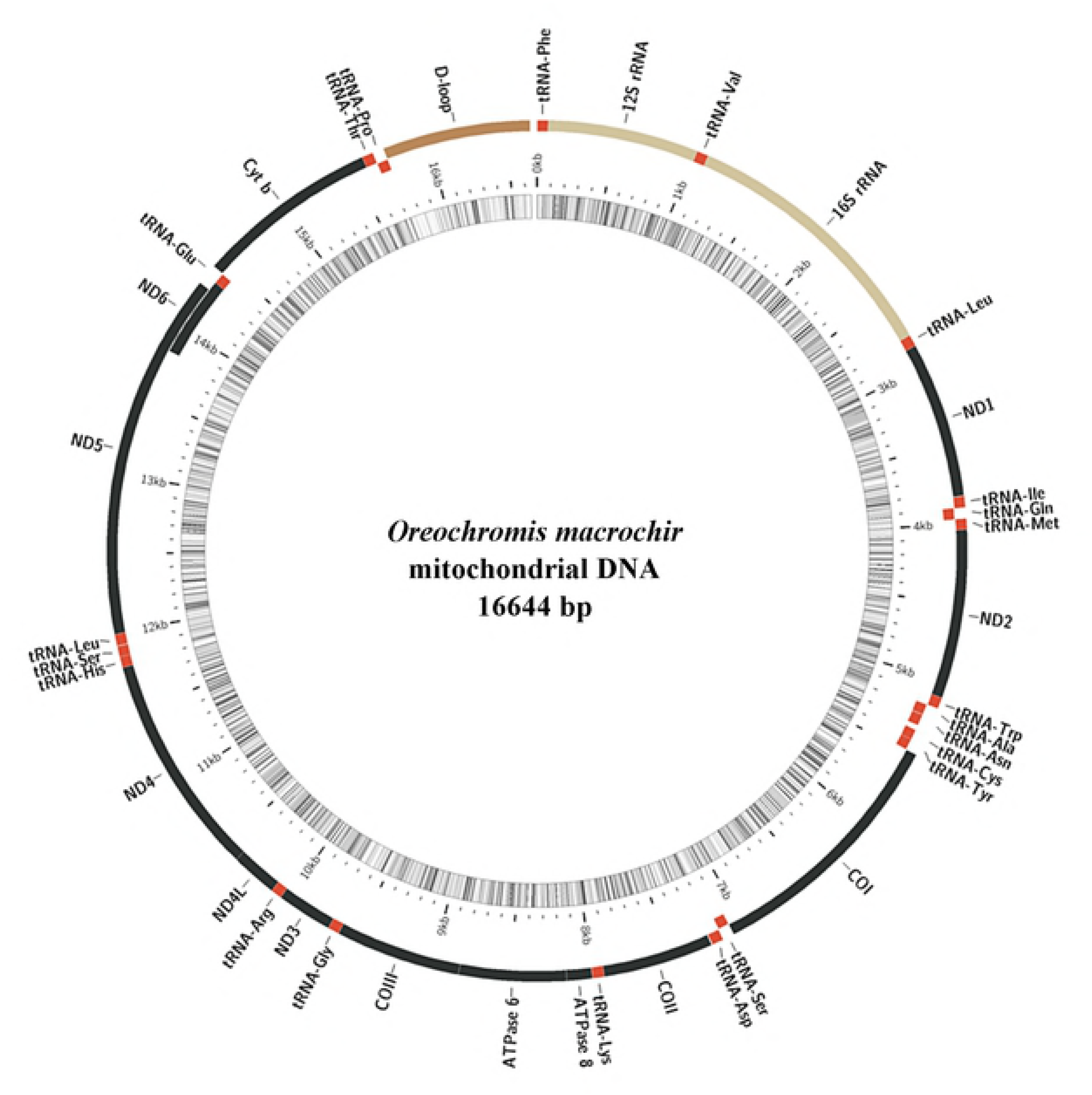
Mitogenomc organisation of *O. macrochir* generated by MitoAnnotator. The genes on the outer side of the circle are coded on the H-strand while those on the inner circle arc coded with L-strand. *(). andersonii* showed similar organization.

The overall nucleotide base composition was very similar between the two species. For *O. andersonii*; T=26.3%, C=30.1%, A=28.0%, G=15.6 % whereas *O. macrochir* T=26.2%, C=30.2%, A=28.1%, G=15.5%. Further, the GC% and AT% were the same for both species at 45.7 and 54.3 respectively (Table 3). This base composition is very similar to what has been reported for other *Oreochromis* species [30,32] and other species as well [31, 32]. The lower value for GC compared to AT is another common feature which has been observed in most vertebrate mitogenome resulting from anti-bias against G in the third codon position. The species in this study had lower values of G in all the three codon positions of protein coding genes and non-coding control region (Table 3). Furthermore, the GC-skew and AT-skew which describe the overall patterns of nucleotide composition in DNA sequences [40], were -0.317 and 0.031 for *O. andersonii* and -0.322 and 0.035 for *O. macrochir* respectively. This result shows an excess of G over C and excess of A over T (Table 3).

**Table 3.** Base percent composition of complete mitogenome sequences of *O. andersonii* (16642 bp) and *O. macrochir* (16644 bp) generated in MEGA 7.0.26

| Codon position | % Nucleotide composition | Complete mitogenome |  | Protein coding genes |  | ND6 |  | tRNAs |  | rRNAs |  | Control Region |  |
| --- | --- | --- | --- | --- | --- | --- | --- | --- | --- | --- | --- | --- | --- |
|  |  | * *O.A* | * *O.M* | *O.A* | *O.M* | *O.A* | *O.M* | *O.A* | *O.M* | *O.A* | *O.M* | *O.A* | *O.M* |
| 1 | T | 27.0 | 27.0 | 31.0 | 31.0 | 35.0 | 35.0 | 26.0 | 26.0 | 19.0 | 19.0 | 32.0 | 31.0 |
|  | C | 31.3 | 31.3 | 30.1 | 30.1 | 8.0 | 8.0 | 22.8 | 23.0 | 26.6 | 26.7 | 20.4 | 19.1 |
|  | A | 27.7 | 27.9 | 25.3 | 25.2 | 6.9 | 6.9 | 27.2 | 27.2 | 32.4 | 32.6 | 32.0 | 34.3 |
|  | G | 13.8 | 13.8 | 13.9 | 14.0 | 50.0 | 50.0 | 23.6 | 23.4 | 21.7 | 21.7 | 15.5 | 15.5 |
| 2 | T | 24.0 | 24.0 | 27.0 | 27.0 | 43.0 | 43.0 | 31.0 | 31.0 | 21.0 | 22.0 | 31.0 | 30.0 |
|  | C | 29.8 | 30.0 | 36.2 | 36.2 | 21.8 | 22.4 | 20.7 | 20.7 | 25.7 | 25.6 | 20.5 | 23.6 |
|  | A | 29.3 | 29.3 | 26.6 | 26.8 | 10.9 | 10.9 | 29.0 | 29.0 | 32.5 | 32.7 | 32.5 | 32.7 |
|  | G | 16.5 | 16.4 | 10.0 | 9.9 | 24.1 | 24.1 | 19.9 | 19.9 | 20.4 | 20.3 | 15.6 | 13.6 |
| 3 | T | 27.0 | 27.0 | 24.0 | 24.0 | 37.0 | 37.0 | 24.0 | 24.0 | 21.0 | 21.0 | 35.0 | 38.0 |
|  | C | 29.3 | 29.2 | 30.8 | 31.0 | 1.1 | 1.1 | 21.2 | 21.0 | 28.0 | 28.2 | 20.8 | 19.4 |
|  | A | 27.0 | 26.9 | 26.9 | 26.8 | 20.5 | 19.4 | 28.4 | 28.6 | 31.9 | 31.9 | 31.2 | 29.4 |
|  | G | 16.3 | 16.3 | 17.9 | 17.9 | 41.4 | 42.5 | 26.4 | 26.3 | 19.5 | 19.2 | 13.0 | 13.3 |
| Total composition | T(U) | 26.3 | 26.2 | 27.5 | 27.4 | 38.3 | 38.3 | 27.0 | 27.0 | 20.5 | 20.4 | 32.9 | 33.0 |
|  | C | 30.1 | 30.2 | 32.4 | 32.5 | 10.3 | 10.5 | 21.6 | 21.6 | 26.8 | 26.8 | 20.5 | 20.7 |
|  | A | 28.0 | 28.1 | 26.2 | 26.2 | 12.8 | 12.3 | 28.2 | 28.2 | 32.3 | 32.4 | 31.9 | 32.1 |
|  | G | 15.6 | 15.5 | 13.9 | 13.9 | 38.5 | 38.9 | 23.2 | 23.3 | 20.5 | 20.4 | 14.7 | 14.1 |
|  | % GC | 45.7 | 45.7 | 46.3 | 46.4 | 48.9 | 49.4 | 44.7 | 44.9 | 47.3 | 47.2 | 35.2 | 34.8 |
|  | % AT | 54.3 | 54.3 | 53.7 | 53.6 | 51.1 | 50.6 | 55.3 | 55.1 | 52.7 | 52.8 | 64.8 | 65.2 |
|  | GC Skew | -0.317 | -0.322 | -0.400 | -0.401 | 0.578 | 0.575 | 0.036 | 0.038 | -0.133 | -0.136 | -0.165 | -0.190 |
|  | AT Skew | 0.031 | 0.035 | -0.024 | -0.022 | -0.499 | -0.514 | 0.022 | 0.022 | 0.223 | 0.227 | -0.015 | -0.014 |
|  | Total length (bp) | 16642 | 16644 | 11427 | 11427 | 522 | 522 | 1554 | 1554 | 2638 | 2638 | 925 | 927 |

\**O.A*= *Oreochromis andersonii*
\**O.M*= *Oreochromis macrochir*

Intergenic overlaps were observed in 4 genes and spacers in 10 in the mitogenome of *O. andersonii* and *Oreochromis macrochir*. The most significant overlap of 461 nucleotides was observed between *ND5* and *ND6*. This large overlap is not a common feature among most reported teleosts mitogenomes. Overlap of 10 nucleotides between *ATPase 8* and *ATPase 6* was also observed for both species followed by *ND4L* and *ND4* gene with overlap of 7 nucleotides. Intergenic spacers totalled 56 bp in 10 regions for both *O. andersonii* and *O. macrochir* (see Table 2). The most significant intergenic spacers in both species were between *tRNA^Asn^* and *tRNA^Cys^* (33 nucleotides) followed by *tRNA^Asp^* and *COII* (5 nucleotides). Again intergenic overlaps and spacers follow what has been reported for most vertebrate mitogenomes inclusive of different fish species except for the overlap between *ND5* and *ND6*.

### Protein coding genes

The nucleotide composition of the protein coding genes was very similar between *O. andersonii* and *O. macrochir* (Table 3). The total GC% composition was 46.3 and AT% was 53.7 for *O. andersonii* and 46.4 and 53.6 for *O. macrochir*. The GC-skew for *O. andersonii* and *O. macrochir* were -0.400 and -0.401 and the AT-skew values were -0.024 and -0.022 respectively. However, the L-strand gene *ND6* had a positive GC-skew value for both species (Table 3). Again it showed overall anti-G bias supporting earlier findings in fish mitogenomes [26, 27, 33]. However, unlike many authors who report strong anti-G bias on third codon positions, both *O. andersonii* and *O. macrochir* were observed to have the strong anti-G bias on the second codon position (O. *andersonii* = 10.0, *O. macrochir* = 9.9).

The adaptive radiation of African cichlids is unparalleled so far among the vertebrates. To understand the role of mitogenome protein coding genes in the evolution of these cichlid species we analysed the rate of non synonymous (*dN*) and synonymous (*dS*) nucleotide substitutions in 13 protein coding genes of 20 species. These included; Riverine, Lake Victoria, Lake Tanganyika and Lake Malawi species *(Sarotherodon melanotheron, Coptodon zillii, Oreochromis aureus, O. niloticus, O. niloticus* GIFT, *O. mossambicus, O. variabilis, O. andersonii, O. macrochir, O. esculentus, Oreochromis* sp ‘red tilapia’, *Petrochromis trewavasae, Tropheus duboisi, Astatotilapia calliptera, Cynotilapia afra*, *Maylandia zebra, Pundamilia nyererei, Tylochromis polylepis, Haplochromis burtoni, Lethrinops lethrinus)*. The mean value of *dN* and *dN/dS* was highest in *ND6* gene and lowest in *COIII* gene for all the 13 protein coding genes (Fig 2). Our findings indicate that all protein coding genes evolved under purifying selection except for *ND6* gene which had an elevated rate of *dN/dS* indicative of evolution under positive selection. We can conclude that *ND6* gene may have evolved more rapidly than any other protein coding gene among these African cichlids.

**Fig 2.**
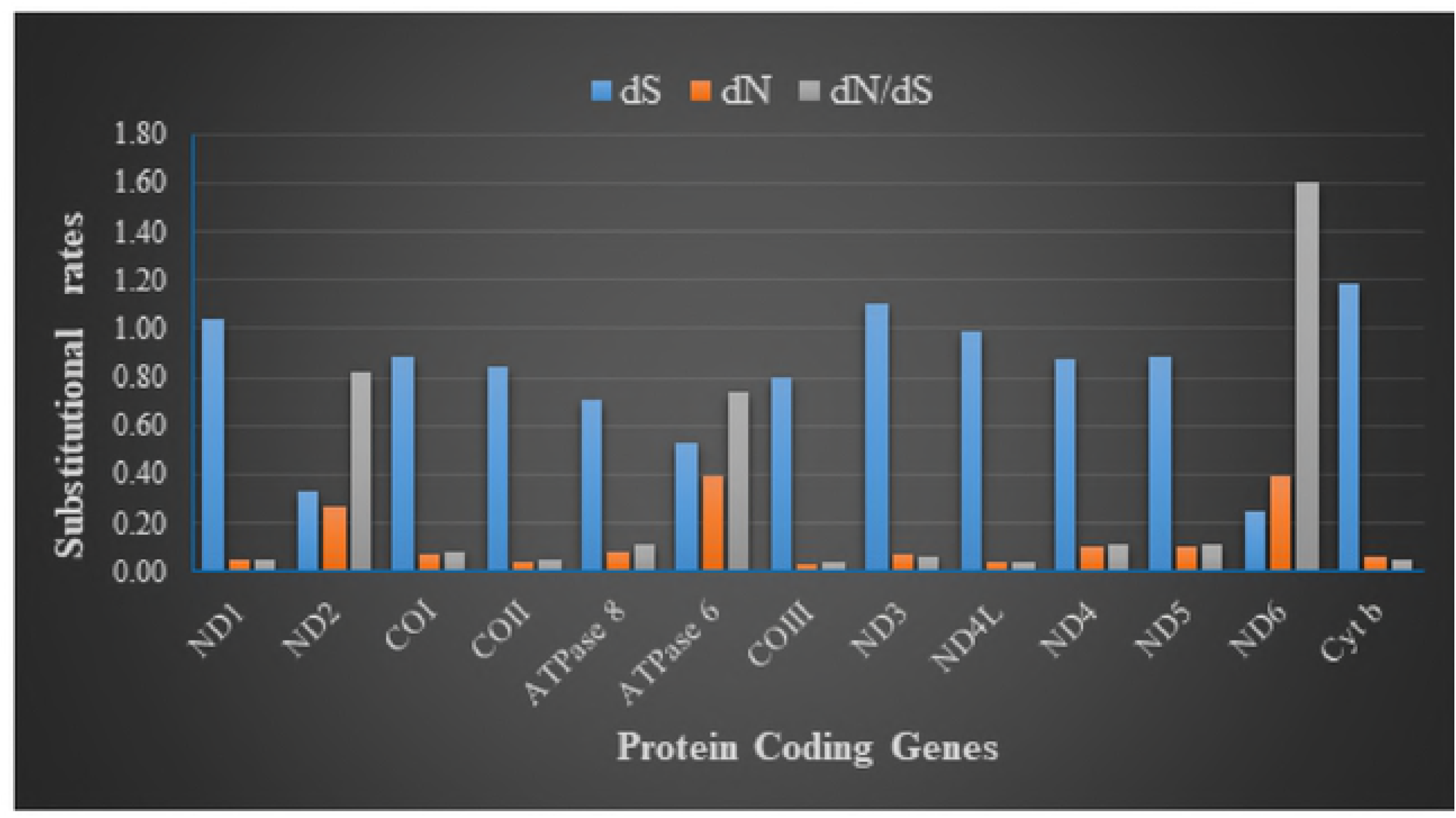
The rate of non-synonymous substitution (c/5), rate of synonymous substitution *(dN)* and the ratio of c/5 and *dN* for each protein coding gene.

The overall *p*-genetic distance was used to measure the conservation of the protein coding genes in the mitogenome of these 20 cichlid species. Calculation was performed on the 1^st^ and 2^nd^, 3^rd^ and whole sequence codon positions. On the 1^st^ and 2^nd^ codon positions, the highest overall mean p-distance was on gene *ND6* (0.1581) followed by *ATPase* 6 (0.1268) and least was gene *COIII* (0.0193). For the whole sequence, the highest overall *p*-distance was recorded in *ATPase* 6 (0.1389) followed by *ND6* (0.1288) and *ATPase* 8 (0.0990) had the least value (Fig 3). Based on these results *ND6* likely may have the highest evolutionary rate and *COIII* been the most conserved gene among the mitogenome protein coding genes of these cichlid species which could have radiated into many species due to geographical isolation [1].

**Fig. 3.**
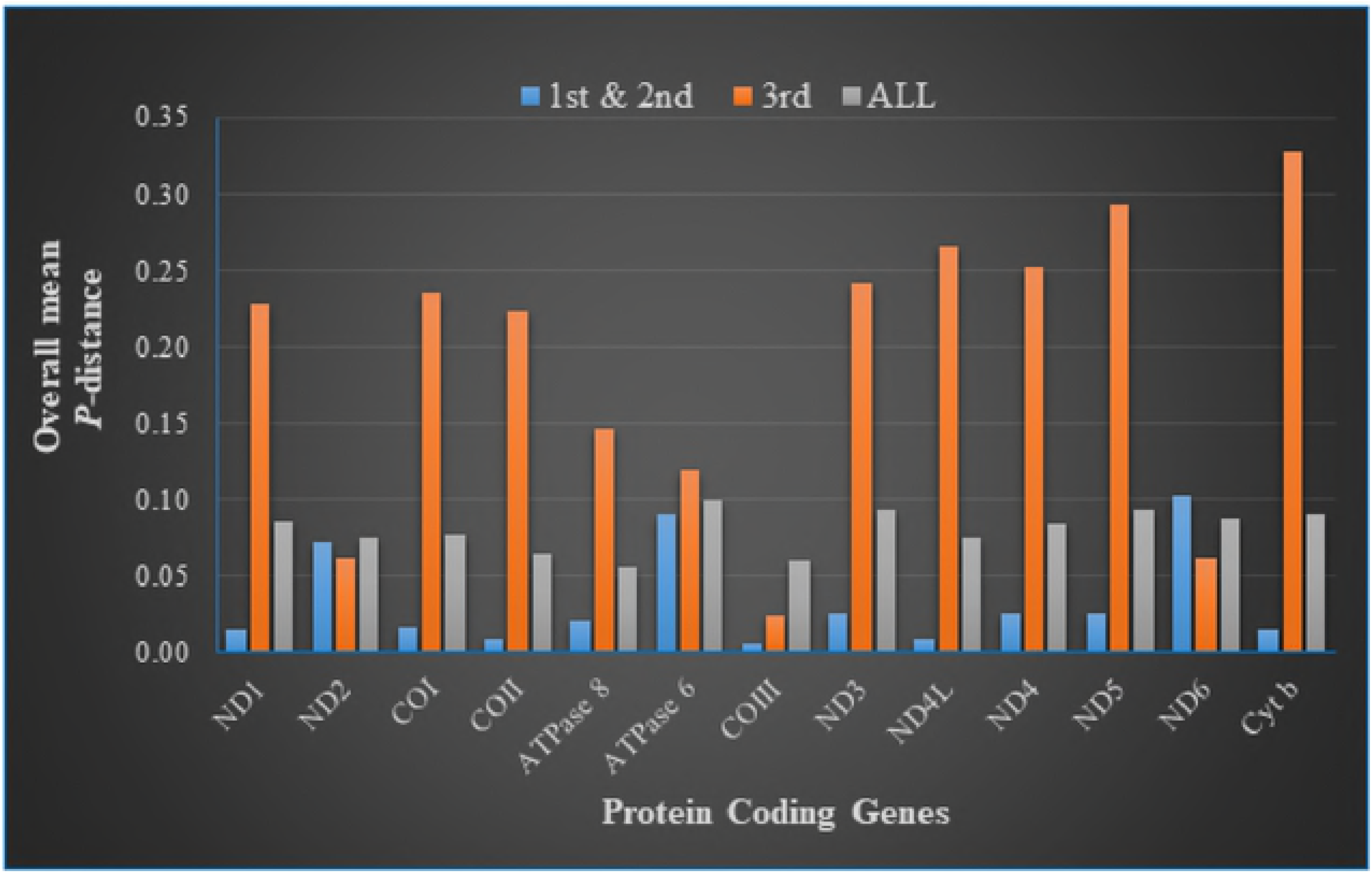
Estimates of average evolutionary divergence over all sequence pairs of 20 cichlid species for each of the 13 protein coding genes calculated based on codon positions Γ’ + 2^nd^, 3^rd^ and full sequence

### Ribosomal and Transfer RNAs

The *tRNA* genes for both *O. andersonii* and *O. macrochir* possessed anti-codons that match the mitochondrial vertebrate code. The length of the *tRNAs* for both species ranged between 66 -74 bp while the total length was 1554 bp for both *O. andersonii* and *O. macrochir*. For both species all the 22 *tRNAs* with exception of *tRNA^Ser(GCT)^* inferred secondary structures folded into classic cloverleaf. The common features of the secondary structures were: a 7 bp aminoacyl stem and anticodon loop, 5 bp TΨC and anticodon arms, and 4 bp DHU arm (S1 Fig). Similar results have been reported on South American catfish and Asian arowana [26, 33]. However, non-complementary pairing and size variations in the secondary structures were observed in both species.

The two *rRNAs* for both species had a total length of 944 bp and 1694 bp which was within the range reported for vertebrate mitogenomes. The nucleotides percent compositions of *rRNAs* of *O*. *andersonii* (T=20.5, C= 26.8, A=32.3, G=20.5) and *O. macrochir* (T= 20.4, C= 26.8, A=32.4, G=20.4) 1 1 were very similar. They also showed a higher percentage of AT than GC pairs (52.7, 47.3 for *O. andersonii*; 52.8, 47.2 for *O. macrochir*) (Table 3)

### Non-coding region

The mitochondrial non-coding region or control region of *O. andersonii* and *O. macrochir* was located between *tRNA^pro^* and *tRNA^phe^* typical for vertebrate mitogenomes. Its length was 925 bp for *O. andersonii* and 927 bp for *O. macrochir*. The nucleotide base composition was similar between the two species although more variable compared to other regions of the mitogenome (Table 3). The AT% (35.2:34.8, *O. andersonii: O. macrochir*) composition to GC% (64.8:65.2, *O. andersonii: O. macrochir*) was also more highly skewed towards AT compared to other regions of the mitogenome due to anti-G bias on the third codon position.

*Oreochromis andersonii* and *O. macrochir* control regions were aligned with four other species from the same genus to characterize the control region. Domain 1 consisted of a hypervariable region with a length of 281 bp for both species including a Termination -associated Sequence (TAS) with motif-ATGCAT similar to a putative TAS of *O. aureus* [30]. Domain II or central conserved region was identified with three conservative sequence blocks (CSB) which are involved in heavy-H strand replication [58]. The first was CSB-F (ATGTAGTAAGAGCCCACC) followed by CSB-E (AAGGACAGTACTTGTGGGGGT) and then CBS-D (TATTCCTGGCATCTGGTTCCT) to complete domain II for both species. The third domain at the 3 end of the control region consisted of CBS-1 *(O. andersonii -* ATTACATAACTGATATCAAGAGCATA; *O. macrochir-* ACCACATAACTGATATCTAGAGCATA) and CBS-2 (AAACCCCCCCTACCCCC). The last conservative block observed was CBS-3 (TGCAAACCCCCCGGAAACAG) (Fig 4)

**Fig 4.**
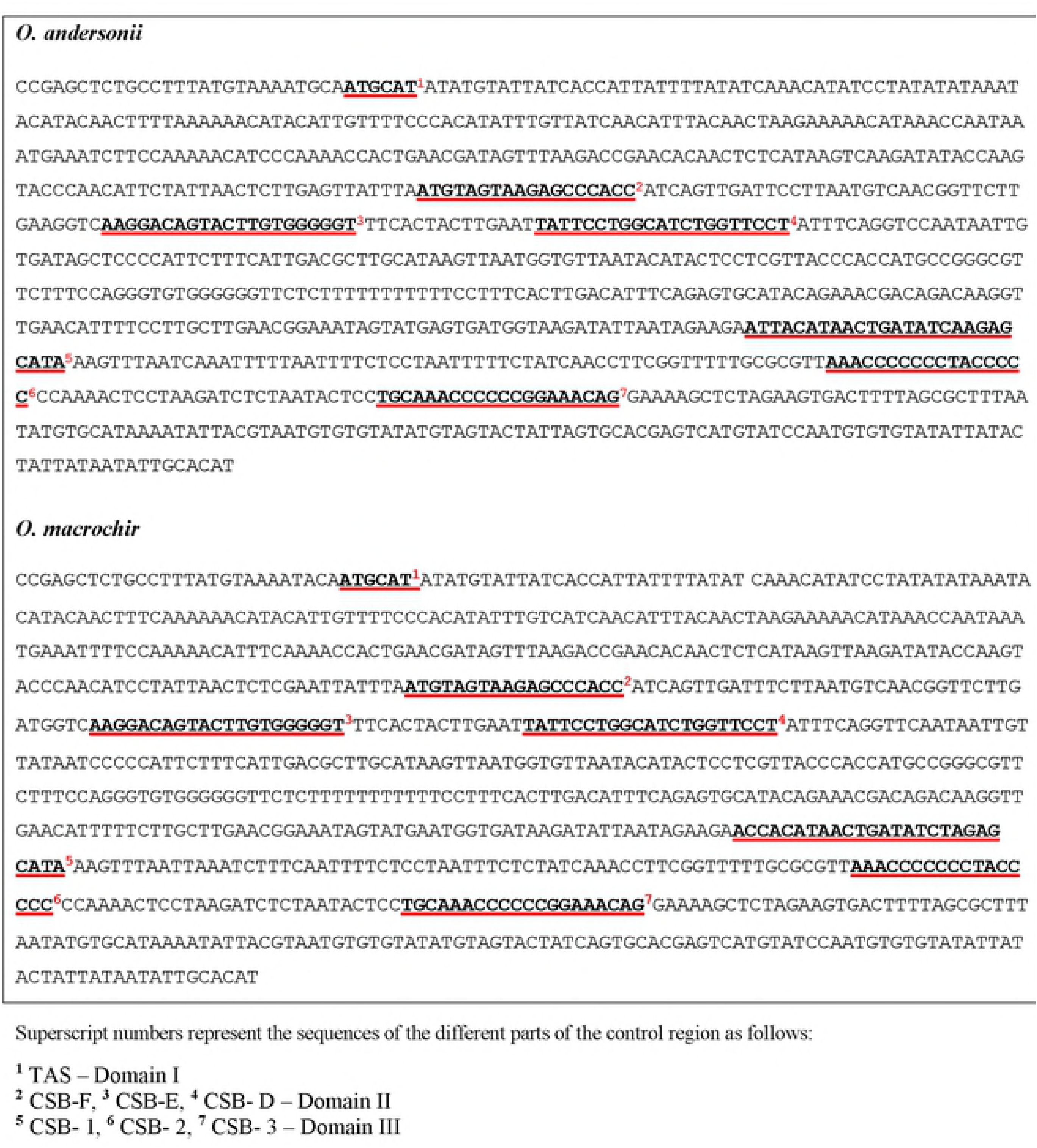
Different parts of the control region of *O. andersonii* and *O. macrochir*

### Amino acid composition

For both *O. andersonii* and *O. macrochir* leucine, proline, serine, threonine, asparagine were the most frequently translated amino acids from the mitochondrial genome (Table 5 and Fig 5). This is similar to the findings on other fish species [26, 33]. The relative synonymous codon usage (RSCU) indicated that the most frequently used codon in the mitogenome of the two species was GCC for alanine with *O. andersonii* having an RSCU value of 1.74 while *O. macrochir* had 1.67. The second highest codon usage values were for serine (UCU) for *O. andersonii* having RSCU=1.54 and Arginine (CGC) for *O. macrochir* having an RSCU=1.58. RSCU values of 0.40 and 0.43 were observed to be the least for alanine (GCG) for *O. andersonii* and *macrochir* respectively.

**Fig 5.**
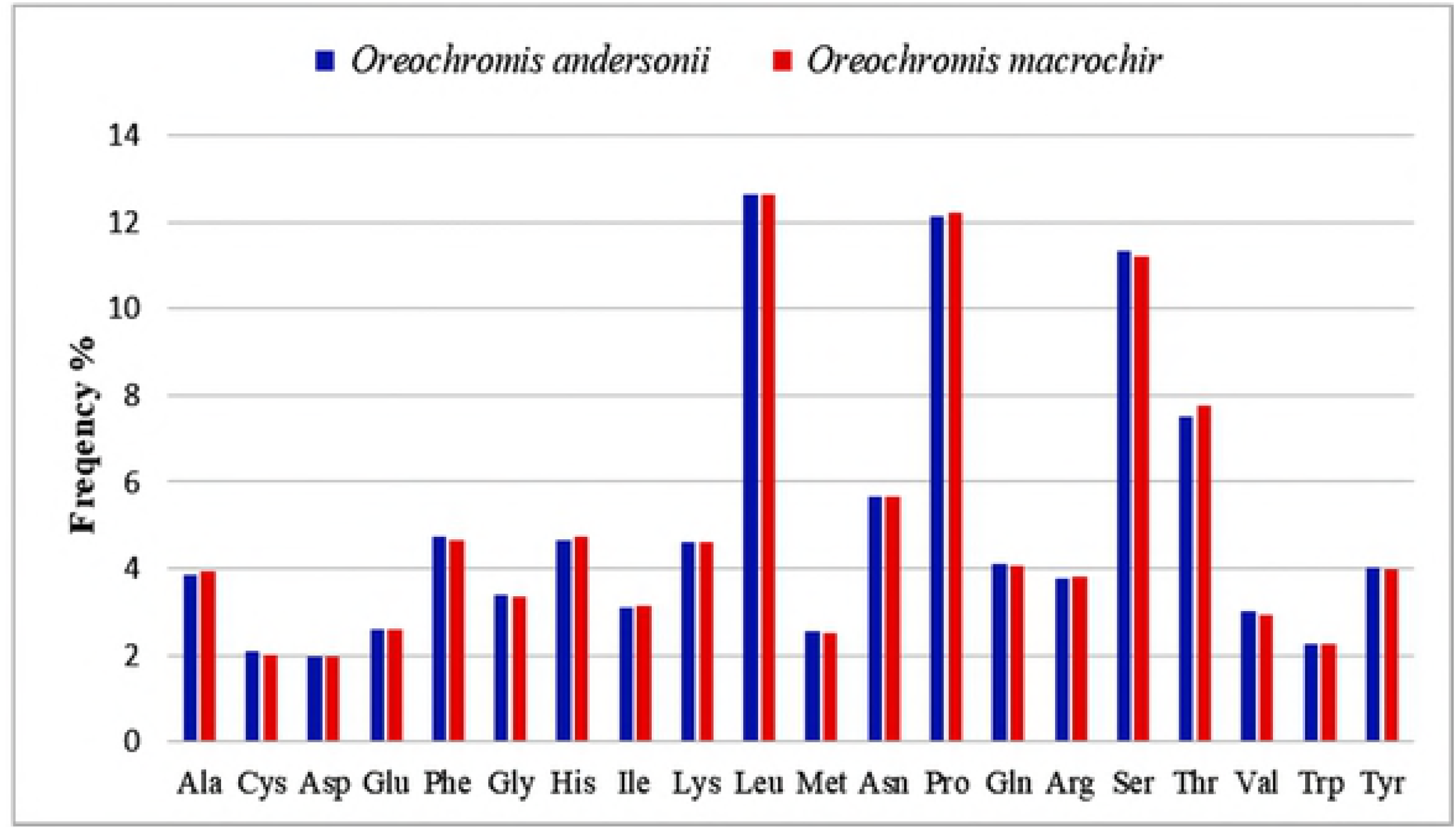
Amino acid frequency in the complete mitogenome of *O. andersonii* and *O. macrochir*

### Phylogenetic analysis

Phylogenetic analysis using 12 protein coding genes of 29 cichlid species and 18 other species belonging to 7 different families of mainly Southern African fish were used to examine the phylogenetic placement of *O. andersonii* and *O. macrochir* among the cichlids of Africa. Most of the families were monophyletic. The consensus tree forms a clear clade consisting of genera *Oreochromis* (maternal mouthbrooders), *Sarotherodon* (biparental and paternal mouthbrooders) and *Coptodon* (substrate spawners) agreeing with the classification of Trewavas of these species [2]. This classification seems to indicate that the development of the mouthbrooding reproductive behaviour emerged later after the substrate and biparental. This phylogenetic relationship was also observed by Nagal [59]. The close relatedness of *Sarotherodon melanotheron* to *Oreochromis aureus* rather forming a monophyletic clade may suggest a need to redefine the genus [32, 59].

The three *Oreochromis* species (*O. macrochir*, *O. andersonii* and *O. mossambicus*) from southern Africa in this classification seem to have evolved later compared to the west and east African species. Data from the consensus tree in this study placed *O. andersonii* closely related to *O. mossambicus* (Fig 6). This is in agreement with other traditional classifications [20, 60]. The close relatedness of these two species though not sharing the same habitat may indicate that they may have separated most recently. On the other hand, *O. macrochir* clusters with both *O. mossambicus* and *O. andersonii* indicative of a closer phylogenetic relationship among these species of Southern Africa. The consensus tree for the other Lake Malawi cichlids is similar to what has been reported by [44]. The families Mugilidae, Centrarchidae and Anabantidae were the most closely related to the cichlids (Fig 6).

**Fig 6.**
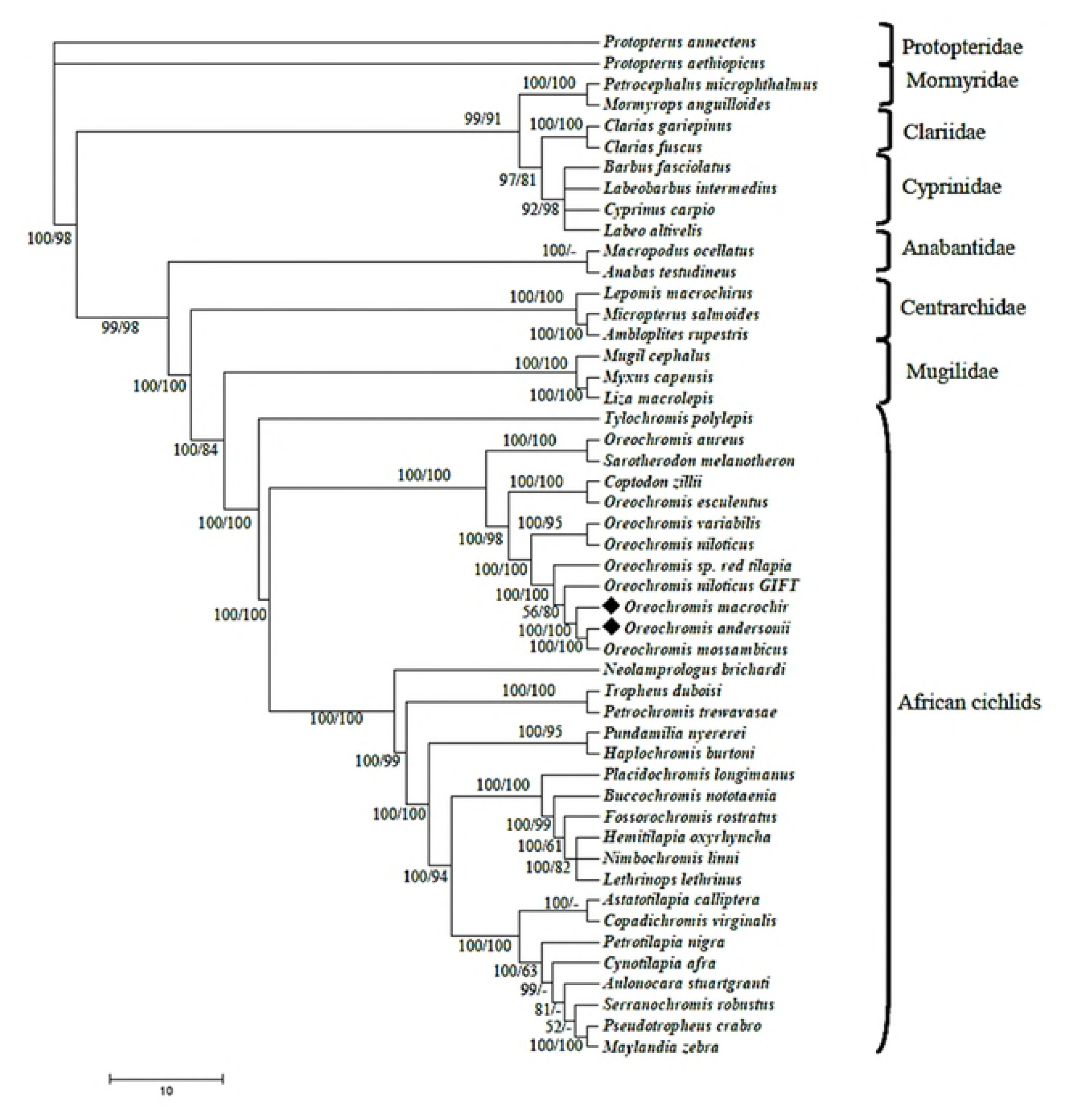
The phylogenetic tree showing the position of *O. andersonii* and *O. macrochir* with other 47 species based on Bayesian posterior probabilities and bootstrap values of ML (numbers in branches) of 12 concatenated protein genes (without *ND6).*

## Conclusion

This study has determined the complete mitochondrial DNA sequences of *O. andersonii* (16623 bp) and *O. macrochir* (16624 bp) and further characterization has confirmed their similarity to teleost vertebrate mitochondrial. Using Bayesian posterior probabilities and Maximum Livelihood analysis of concatenated mitochondrial genome of 12 protein coding genes, consensus trees has revealed a close phylogenetic relationship between *O. andersonii* and *O. mossambicus*. Further, *O. macrochir* was found to be closely related to both *O. andersonii* and *O. mossambicus*. The protein coding genes evolved under purifying selection except for *ND6* which indicated evolution under positive selection. Gene *ND6* likely may have the highest evolutionary rate and *COIII* been the most conserved gene. More work is needed to characterize the southern Africa cichlids as they are important species especially for capture fisheries, aquaculture development and understanding biogeographic history of African cichlids.

## Acknowledgments

We thank the Department of Fisheries – Zambia staff for support during sample collection.

## Supporting information

S1 Fig. Secondary structures of 22 *tRNAs* of mitochondrial genome of *O. andersonii* (A) and *O. macrochir* (B) generated by tRNAScan-SE 2.0

S1 Table. Designed primers used for amplification and sequencing of complete mitogenome of *O. andersonii* and *O. macrochir*

S2 Table. Complete automated annotation of *O. andersonii* and *O. macrochir* mitochondrial genome.

## References

1. Brawand D, Wagner CE, Li YI, Malinsky M, Keller I, Fan S, et al. The genomic substrate for adaptive radiation in African cichlid fish. Nature. 2014; 513:375–381. doi :10.1038/nature13726.

2. Trewavas E. Tilapiine fishes of the Genera Sarotherodon, Oreochromis, and Danakilia. London: British Museum (Natural History); (1983).

3. Skelton P. Freshwater Fishes of Southern Africa. Second edition. South Africa: Struik Publishers (Pty) Ltd; 2001.

4. Froese R, Pauly D; 2016 [cited ]. Database: FishBase [Internet]. Available from: http://www.fishbase.org

5. Lind CE, Brummett RE, Ponzoni RW. Exploitation and conservation of fish genetic resources in Africa: issues and priorities for aquaculture development and research. Reviews in Aquaculture. 2012; 4:125–141.

6. Tweddle D, Marshall BE; 2007. [cited ]. Database: Oreochromis andersonii. The IUCN Red List of Threatened Species [Internet]. Available from: http://dx.doi.org/10.2305/IUCN.UK.2007.RLTS.T60623A12385801.en

7. Tweddle D, Peel RA, Weyl OLF. Challenges in fisheries management in the Zambezi, one of the great rivers of Africa. Fisheries Management and Ecology. 2015; 22:99–111.

8. Canonico GC, Arthington A, McCrary JF, Thiem ML. The effects of introduced tilapias on native biodiversity. Aquatic Conserv: Mar Freshw Ecosyst. 2005; 15:463–483. doi:10.1002/aqc.699.

9. Deines AM, Bbole I, Katongo C, Feder JL, and Lodge DM. Hybridisation between native Oreochromis species and introduced Nile tilapia O. niloticus in the Kafue River, Zambia. Afr J Aquat Sci. 2014; 39:23–34.

10. Bbole I, Katongo C, Deines AM, Shumba O, Lodge DM. Hybridization between non-indigenous Oreochromis niloticus and native Oreochromis species in the lower Kafue River and its potential impacts on fishery. J. Ecol. Nat. Environ. 2015; 6(6):215–225.

11. Crayon-Thomas E. Background history on the research and use of Oreochromis niloticus in Zambia and their use on a private fish farm (Kalimba Farm) between 1985 and 1993. Technical Consultation on Species for Small Reservoir Fisheries and Aquaculture in Southern Africa. Livingstone, Zambia, 7–11 November 1994

12. Kefi AS, Kang’ombe J, Kassam D, Katongo C. Growth, reproduction and sex ratios in Oreochromis andersonii (Castelnau, 1861) fed with varying Levels of 17α-methyl testosterone. J Aquac Res Dev. 2012; 3:130–137.

13. Musuka CG, Musonda FF. Current and future prospects of commercial fish farming in Zambia. Int J Bioflux Soc. 2012 5:79–87.

14. Nsonga A. Indigenous fish species a panacea for cage aquaculture in Zambia: A case for Oreochromis macrochir (Boulenger, 1912) at Kambashi out-grower scheme. Int. j. fish. Aquat. 2014; 2(1):102–105.

15. Kefi AS, Kang’ombe J, Kassam D, Katongo C. Optimal Dietary Plant Based Lipid on Growth of Oreochromis andersonii (Castelnau, 1861). Turk J Fish Aquat Sci. 2013; 13:503–508.

16. Salzburger W, Meyer A, Baric S, Verheyen E, Sturmbauer C. Phylogeny of the Lake Tanganyika Cichlid Species Flock and Its Relationship to the Central and East African Haplochromine Cichlid Fish Faunas. Syst Biol. 2002; 51(1):113–135.

17. Salzburger W, Meyer A. The species flocks of East African cichlid fishes: recent advances in molecular phylogenetics and population genetics. Naturwissenschaften. 2004; 91:277–290.

18. Meyer BS, Matschiner M, Salzburger W. A tribal level phylogeny of Lake Tanganyika cichlid fishes based on a genomic multi-marker approach. Mol Phylogenet Evol. 2015; 83:56–71.

19. Kocher TD, Lee W, Sobolewska H, Penman D, Andrew B. A genetic linkage map of a cichlid fish the tilapia (Oreochromis niloticus). Genetics. 1998; 148:1225–1232.

20. Sodsuk PK, McAndrew BJ, Turner Gf(1995) Evolutionary relationships of the Lake Malawi Oreochromis species: evidence from allozymes. J Fish Biol. 1995; 47:321–333.

21. Feresu-Shonhiwa F, Howard JH. Electrophoretic identification and phylogenetic relationships of indigenous tilapiine species of Zimbabwe. J Fish Biol. 1998; 53:1178–1206.

22. Katongo C, Koblmüller S, Duftner N, Makasa L, Sturmbauer C. Phylogeography and speciation in the Pseudocrenilabrus philander species complex in Zambian Rivers. Hydrobiologia. 2005; 542:221–233. doi: 10.1007/s10750-004-1389-x

23. D’Amato ME, Esterhuyse MM, Van der Waal BCW, Brink D, Volckaert FAM. Hybridization and Phylogeography of the Mozambique tilapia Oreochromis mossambicus in Southern Africa evidenced by mitochondrial and microsatellite DNA genotyping. Conserv. Genet. 2007; 8: 475–488.

24. Katongo C, Koblmüller, S, Duftner N, Mumba L, Sturmbauer C. Evolutionary history and biogeographic affinities of the Serranochromine cichlids in Zambian rivers. Mol.Phylogenetics Evol. 2007; 45:326–338. doi:10.1016/j.ympev.2007.02.011

25. Peng Z, Wang J, He S. The complete mitochondrial genome of the helmet catfish Cranoglanis bouderius (Siluriformes: Cranoglanididae) and the phylogeny of otophysan fishes. Gene. 2006; 376:290–297.

26. Yue GH, Liew WC, Orban L. The complete mitochondrial genome of a basal teleost, the Asian arowana (Scleropages formosus, Osteoglossidae). BMC Genomics. 2006; 7:242.

27. Liu Y, Cui Z. The complete mitochondrial genome sequence of the cutlassfish Trichiurus japonicas (Perciformes:Trichiuridae): Genome characterization and phylogenetic considerations. Mar Genomics. 2009; 2:133–142.

28. Zhang X, Yue B, Jiang W, Song Z. The complete mitochondrial genome of rock carp Procypris rabaudi (Cypriniformes: Cyprinidae) and phylogenetic implications. Mol Biol Rep. 2009; 36:981–991.

29. Cheng YZ, Wang RX, Xu TJ. The mitochondrial genome of the spinyhead croaker Collchthys lucida: Genome organization and phylogenetic consideration. Mar Genomics. 2011; 4:17–23.

30. He A, Luo Y, Yang H, Liu L, Li S, Wang C. Complete mitochondrial DNA sequences of the Nile tilapia (Oreochromis niloticus) and Blue tilapia (Oreochromis aureus): genome characterization and phylogeny applications. Mol Biol Rep. 2011; 38:2015–2021. DOI 10.1007/s11033-010-0324-7

31. Wang C, Wang J, Yang J, Song X, Chen Q, Xu J, Yang Q, Li S. Complete mitogenome sequence of black carp (Mylopharyngodon piceus) and its use for molecular phylogeny of leuciscine fishes. Mol Biol Rep. 2012; 39:6337–6342. doi: 10.1007/s11033-012-1455-9

32. Kinaro ZO, Xue L, Volatiana JA. Complete mitochondrial DNA sequences of the Victoria tilapia (Oreochromis variabilis) and Redbelly Tilapia (Tilapia zilli): genome characterization and phylogeny analysis. Mitochondrial DNA Part A. 2015; 27(4): 2455–2457.

33. Villela LC, Alves AL, Varela ES, Yamagishi ME, Giachetto PF, da Silva NM, et al. Complete mitochondrial genome from South American catfish Pseudoplatystoma reticulatum (Eigenmann & Eigenmann) and its impact in Siluriformes phylogenetic tree. Genetica. 2017. 145:51–66. doi: 10.1007/s10709-016-9945-7.

34. Hall TA. BioEdit: a user-friendly biological sequence alignment editor and analysis program for Windows 95/98/NT. Nucl Acids Symp Ser. 1999; 41:95–98.

35. Lowe TM, Eddy SR. tRNAscan-SE: A program for improved detection of transfer RNA genes in genomic sequence. Nucl Acids Res. 1997; 25:955–964.

36. Kumar S, Stecher G,Tamura K (2016) MEGA7: Molecular Evolutionary Genetics Analysis Version 7.0 for Bigger Datasets. Mol Biol Evol. 2016; 33(7):1870–1874. https://doi.org/10.1093/molbev/msw054

37. Wyman SK, Jansen RK, Boore JL. Automatic annotation of organellar genomes with DOGMA. Bioinformatics. 2004; 20(17):3252–5. doi: 10.1093/bioinformatics/bth352

38. Bernt M, Donath A, Jühling F, Externbrink F, Florentz C, Fritzsch G, et al. MITOS: improved de novo metazoan mitochondrial genome annotation. Mol Phylogenet Evol. 2013; 69(2):313–9.doi: 10.1016/j.ympev.2012.08.023

39. Iwasaki W, Fukunaga T, Isagozawa R, Yamada K, Maeda Y, Satoh TP, et al. MitoFish and MitoAnnotator: a mitochondrial genome database of fish with an accurate and automatic annotation pipeline. Mol Biol Evol. 2013; 30:2531–2540.

40. Perna NT, Kocher TD. Patterns of nucleotide composition at fourfold degenerate sites of animal mitochondrial genomes. J Mol Evol. 1995; 41:353–358. doi: 10.1007/BF01215182.

41. Miya M, Nishida M. The mitogenomic contributions to molecular phylogenetics and evolution of fishes: a 15-year retrospect. Ichthyol Res. 2015; 62:29–71. doi: 10.1007/sl0228-014-0440-9.

42. Edgar RC. MUSCLE: multiple sequence alignment with high accuracy and high throughput. Nucleic Acids Res. 2004; 32:1792–1797.

43. He A-Y, Tang S-J, Jiang Y-T, Li S-F, Wang C-H. Complete mitochondrial genome of blackchin tilapia Sarotherodon melanotheron (Perciformes, Cichlidae). Mitochondrial DNA. 2011; (5-6):171–3. doi: 10.3109/19401736.2011.636439

44. Hulsey CD, Keck BP, Alamillo H, O’Meara BC. Mitochondrial genome primers for Lake Malawi cichlids. Molec Ecol Res. 2013b; 13:347–353.

45. Azuma Y, Kumazawa Y, Miya M, Mabuchi K, Nishida M. Mitogenomic evaluation of the historical biogeography of cichlids toward reliable dating of teleostean divergences. BMC Evol Biol. 2008; 8:215. doi: 10.1186/1471-2148-8-215

46. Hu XX, Lu J, He SY, Li XF, Liu HJ, Han LJ. The complete mitochondrial genome of Astatotilapia burtoni. Mitochondrial DNA. 2016; 27(4):2379–80. doi:10.3109/19401736.2015.1028039

47. Chen L, Song X, Chen X, Dang X, Wang W. The complete mitochondrial genome of the Pundamilia nyererei (Perciformes, Cichlidae). Mitochondrial DNA. 2016; 27(5):3567–8. doi:10.3109/19401736.2015.1074221

48. Qi DS, Zhang SC, Jiang J, Zhang LQ, Fu YY, Du ZM, et al. The complete mitochondrial genome of the Fossorochromis rostratus. Mitochondrial DNA. 2016; 27(6):4284–4285.

49. Zhao J, Gao J. Complete mitochondrial genome of Aulonocara stuartgranti (Flavescent peacock cichlid). Mitochondrial DNA A DNA Mapp Seq Anal. 2017; 28(2):279–280. doi:10.3109/19401736.2015.1118079

50. Mabuchi K, Miya M, Azuma Y, Nishida M. Independent evolution of the specialized pharyngeal jaw apparatus in cichlid and labrid fishes. BMC Evol Biol. 2007; 7:10. doi: 10.1186/1471-2148-7-10.

51. Fischer C, Koblmuller S, Gully C, Schlotterer C, Sturmbauer C, Thallinger GG. Complete mitochondrial DNA sequences of the threadfin cichlid (Petrochromis trewavasae) and the blunthead cichlid (Tropheus moorii) and patterns of mitochondrial genome evolution in cichlid fishes. PLoS One. 2013; 8(6):e67048. doi: 10.1371/journal.pone.0067048.

52. Han C, Li Q, Xu J, Li X, Huang J. Characterization of Clarias gariepinus mitochondrial genome sequence and a comparative analysis with other catfishes. Biologia. 2016;70(9): 1245–1253.

53. Broughton RE, Reneau PC. Spatial covariation of mutation and nonsynonymous substitution rates in vertebrate mitochondrial genomes. Mol Biol Evol. 2006; 23 (8):1516–1524

54. Lavoue S, Miya M, Arnegard ME, Sullivan JP, Hopkins CD, Nishida M. Comparable ages for the independent origins of electrogenesis in African and South American weakly electric fishes. PLoS ONE. 2012; 7(5): E36287.

55. Inoue JG, Miya M, Lam K, Tay BH, Danks JA, Bell J, et al. Evolutionary origin and phylogeny of the modern holocephalans (Chondrichthyes: Chimaeriformes): a mitogenomic perspective. Mol Biol Evol. 2010; 27(11):2576–2586. doi: 10.1093/molbev/msq147

56. Miya M, Kawaguchi A, Nishida M. Mitogenomic exploration of higher teleostean phylogenies: a case study for moderate-scale evolutionary genomics with 38 newly determined complete mitochondrial DNA sequences. Mol Biol Evol. 2001; 18(11):1993–2009.

57. Ronquist F, Huelsenbeck JP. MrBayes 3: Bayesian phylogenetic inference under mixed models. Bioinformatics. 2003; 19:1572–1574.

58. Faber JE, Stepien CA. Tandemly repeated sequences in mitochondrial DNA control region and phylogeography of the Pike-Perches Stizostedion. Mol. Phylogenetics Evol. 1998; 10(3):310–322

59. Nagl S, Tichy H, Mayer WE, Samonte IE, McAnderw BJ, Klein J. Classification and phylogenetic relationships of African tilapiine fishes inferred from mitochondrial DNA sequences. Mol. Phylogenetics Evol. 2001; 20(3):361–374.

60. McAndrew BJ, Majumdar K C. Evolutionary relationships within three Tilapiine genera (Pisces: Cichlidae). Zool J Linnean Soc. 1984; 80:421–435.

